# Lysosomal stress induces amyloid-β aggregate release and reactive transformation in human astrocytes

**DOI:** 10.1101/2025.05.27.656416

**Authors:** Sian Goldsworthy, Emre Fertan, Patricia Dyal, Sarah Domoney, Frederick J. Livesey, David Klenerman, Rickie Patani, Christy Hung

**Affiliations:** UCL Great Ormond Street Institute of Child Health, Zayed Centre for Research into Rare Disease in Children, 20 Guilford Street, London WC1N 1DZ, UK; Yusuf Hamied Department of Chemistry, University of Cambridge Cambridge CB2 1EW, UK; Talisman Therapeutics, Babraham Research Campus, Cambridge, CB22 3AT, UK; UK Dementia Research Institute at University of Cambridge Cambridge CB2 0XY, UK; Human Stem Cells and Neurodegeneration Laboratory, The Francis Crick Institute; 1 Midland Road, London NW1 1AT, UK; Department of Neuromuscular Diseases, Queen Square Institute of Neurology, University College London; London WC1N 3BG, UK; Department of Neuroscience, City University of Hong Kong, 31 To Yuen Street, Kowloon, Hong Kong SAR 999077

## Abstract

Astrocytes are essential for brain homeostasis and are involved in amyloid-β (Aβ) clearance, but whether they can produce and release Aβ aggregates remains unclear. Using human iPSC-derived astrocytes, we show that astrocytes cell autonomously generate small, diffusible Aβ aggregates under baseline conditions. By combining ultrasensitive single-molecule imaging (DNA-PAINT) and immunoassays, we detect intracellular aggregates and their release into the media. Notably, lysosomal membrane damage induced by L-leucyl-L-leucine methyl ester (LLOMe) significantly increases Aβ aggregate secretion without altering their size or morphology. Transcriptomic analysis and cytokine profiling reveal that lysosomal stress triggers a reactive astrocyte phenotype marked by upregulation of inflammatory genes and secreted cytokines. These findings suggest that astrocytes are not merely passive Aβ scavengers but can actively contribute to extracellular Aβ accumulation under lysosomal stress. Our study highlights astrocytes as active players in Alzheimer’s disease pathology.

## Introduction

Alzheimer’s disease (AD) is characterized by the progressive accumulation of amyloid-β (Aβ) aggregates in the brain, traditionally attributed to neuronal overproduction and impaired clearance mechanisms^1^. While neurons are considered as a major source of Aβ, emerging evidence suggests that glial cells, including astrocytes and oligodendrocytes, may also contribute to Aβ pathology. Recent work has demonstrated that oligodendrocytes in both human tissue and mouse models possess the molecular machinery to produce Aβ and can contribute to early plaque formation and neuronal dysfunction^2–4^. However, whether astrocytes, despite their well-established role in Aβ clearance^5^, can themselves produce and release Aβ aggregates remains unclear.

Astrocytes are the most abundant glial cell type in the central nervous system and play crucial roles in neuronal metabolism, maintenance, synaptic activity, and amyloid-β (Aβ) clearance^6^. They respond variably to a wide range of nervous system insults, including traumatic brain injury, spinal cord injury, stroke, inflammation, and neurodegenerative diseases^7^. Astrocytes are known to undergo deleterious reactive transformation in a range of human neurodegenerative diseases^8^, driven by proinflammatory factors from microglia^9^. Notably, astrocyte-specific deletion of the sulfatase modifying factor 1 (*Sumf1*) gene, which is associated with multiple sulfatase deficiency, a lysosomal storage disease, is sufficient to induce degeneration of cortical neurons *in vivo*^10^. Lysosomal failure also underlies the pathogenesis of numerous congenital neurodegenerative disorders and is an early and progressive feature of AD pathogenesis^11–17^. Whether lysosomal damage can alter the handling of Aβ by astrocytes and shift them from clearing Aβ to potential contributors of pathologenesis remains unknown.

Different forms of cellular stress can induce permeabilization and rupture of lysosomes. To cope with the detrimental effects of lysosomal damage, cells possess mechanisms to either degrade or repair damaged endomembranes^18^. If unchecked, lysosomal damage can result in the leakage of intralysosomal components, such as cathepsins, into the cytoplasm, inducing lysosome-dependent necroptosis^19^.

Although the lysosomal system has been extensively studied in human neurons, the potential consequences of lysosomal membrane damage in glial cells, particularly astrocytes, remain understudied.

Here, we use human induced pluripotent stem cell (iPSC)-derived astrocytes and super-resolution single-molecule imaging to demonstrate that astrocytes produce and secrete Aβ aggregates under basal conditions. We show that lysosomal membrane damage induced by L-leucyl-L-leucine methyl ester (LLOMe) significantly enhances the release of Aβ aggregates, without altering their structural properties. In parallel, transcriptomic and cytokine profiling reveal that lysosomal stress triggers a reactive transformation in astrocytes, marked by pro-inflammatory gene expression and cytokine release. Together, these findings identify astrocytes as an unappreciated source of Aβ aggregates and establish a mechanistic link between lysosomal damage, reactive transformation, and Aβ aggregate release.

## Results

### Human iPSC-derived astrocytes produce and release small diffusible Aβ aggregates

To determine whether human astrocytes can produce and release amyloid-β (Aβ) aggregates, we first used a previously published directed differentiation method to generate highly enriched populations of human iPSC-derived astrocytes^20^ and collected both the conditioned media and cell lysates for analysis. We then used a highly sensitive single-molecule pull-down (SiMPull) assay combined with DNA-PAINT super-resolution imaging to detect and characterize individual Aβ aggregates^21^ (**Figure 1A**). SiMPull relies on dual antibody binding and clustering analysis, requiring at least two closely positioned super-resolved assemblies to confirm the specific detection of soluble aggregates^22–24^. This highly sensitive approach allowed us to visualize aggregates at the nanometer scale and compare their abundance and structural features across compartments.

**Figure 1.**
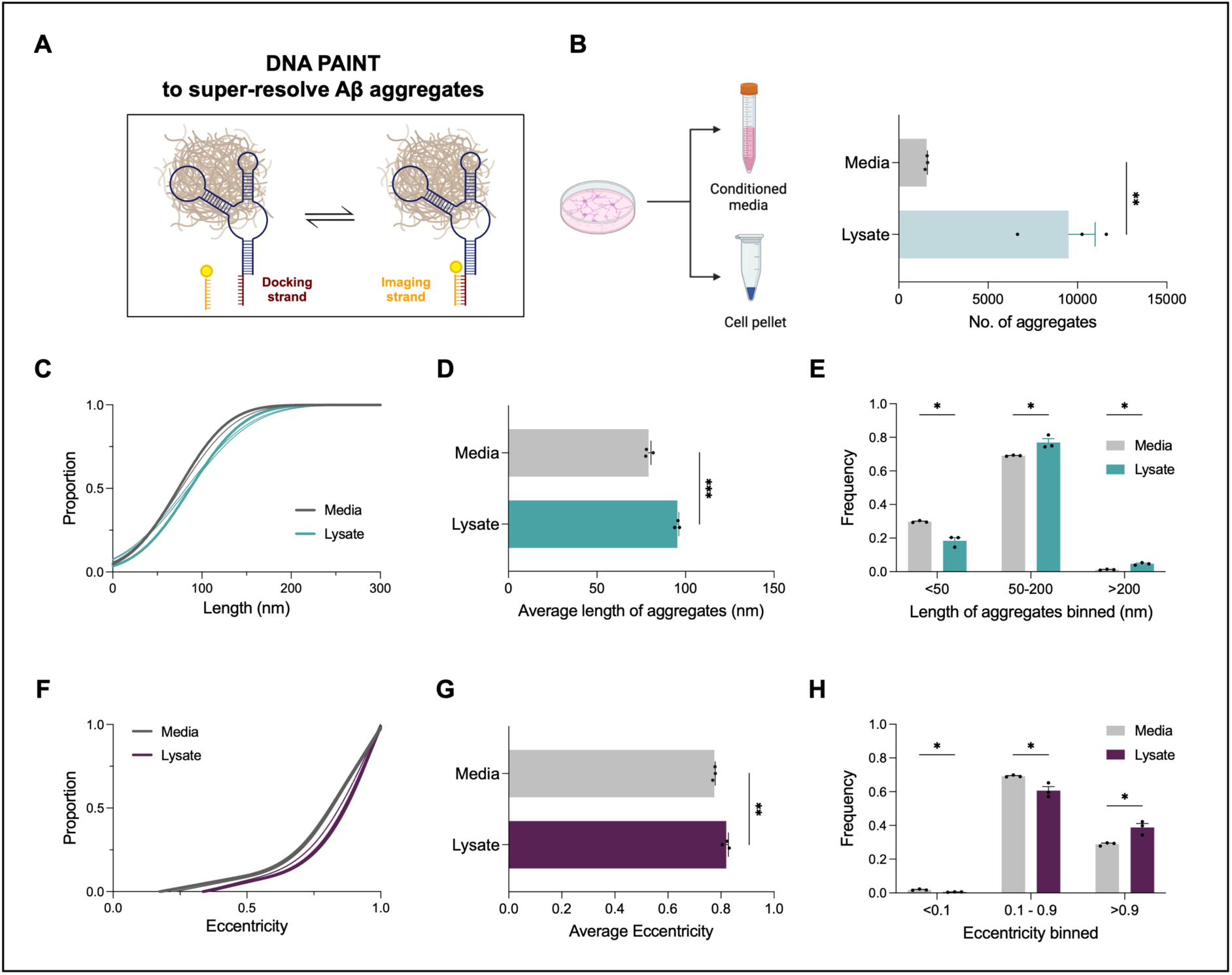
Human iPSC-derived astrocytes produce and release structurally distinct Aβ aggregates. (A) Schematic of DNA-PAINT imaging used to resolve individual Aβ aggregates. (B) Experimental workflow for isolating extracellular (media) and intracellular (lysate) aggregates, and quantification of aggregate numbers using DNA-PAINT. Data represent mean ± SEM from 3 biological replicates. (C) Cumulative frequency plot showing length distribution of Aβ aggregates in media and lysate. (D) Average aggregate length is significantly greater in lysates than in media. Data represent mean ± SEM from 3 biological replicates. (E) Binned length analysis reveals enrichment of small (<50 nm) aggregates in media and longer (>200 nm) aggregates in lysate. (F) Cumulative distribution of aggregate eccentricity indicates more spherical morphology in released aggregates. (G) Average eccentricity is lower in media than in lysate. Data represent mean ± SEM from 3 biological replicates. (H) Frequency of eccentricity categories shows greater proportion of highly eccentric aggregates (>0.9) in lysate. Data represent mean ± SEM from 3 biological replicates. *p < 0.05, **p < 0.01, ***p < 0.001 by unpaired t-test.

Majority of the Aβ aggregates are between 30 nm (resolution limit) and 200 nm in length, consistent with the reported size of synaptotoxic Aβ aggregates from human AD brains^25^. Interestingly, we found a significantly higher number of Aβ aggregates in the cell lysates compared to the media (**Figure 1B**), indicating that astrocytes accumulate Aβ intracellularly but also release a portion into the extracellular space.

Cumulative frequency analysis showed that aggregates in the media were shorter in length compared to those in the lysates (**Figure 1C**), and this was supported by significantly lower average lengths in media samples (**Figure 1D**). We found that small aggregates (<50 nm) were more frequent in the media, while larger aggregates (>200 nm) were enriched in the lysate (**Figure 1E**).

We next examined aggregate shape by analyzing eccentricity, a measure of how elongated or circular the structures are. Aggregates in the media tended to be less elongated, as reflected by lower eccentricity values (**Figure 1F–G**). Lysate samples contained a greater proportion of highly elongated aggregates (eccentricity >0.9), whereas more spherical aggregates (eccentricity <0.2) were found in the media (**Figure 1H**). These data reveal that human astrocytes can produce Aβ aggregates and release a subset into the extracellular space, with secreted aggregates tending to be smaller and less elongated. This supports a model in which astrocytes are active contributors to extracellular Aβ accumulation in the brain.

### Lysosomal membrane damage enhances amyloid-β aggregate release

Having established that human iPSC-derived astrocytes can produce and retain small, diffusible Aβ aggregates intracellularly, we next sought to investigate the cellular mechanisms that might govern their release. Lysosomes play a central role in degrading aggregated proteins, including Aβ, and impaired lysosomal function is increasingly recognized as an early driver of Alzheimer’s disease pathology^11–17^. However, the impact of lysosomal membrane integrity on the intracellular retention or extracellular release of astrocyte-derived Aβ aggregates remains unknown. We therefore induced lysosomal membrane permeabilization using the lysosomotropic agent L-leucyl-L-leucine methyl ester (LLOMe) and examined its consequences on Aβ aggregate distribution and astrocyte state.

To assess lysosomal damage in astrocytes, we performed super-resolution instant structured illumination microscopy (iSIM) imaging of Galectin-3 (Gal3), a cytosolic lectin that binds to exposed glycoproteins on the inner membrane of ruptured lysosomes. Treatment with LLOMe resulted in a significant increase in Gal3-positive puncta, confirming lysosomal membrane damage (**Figure 2A and 2B**).

**Figure 2.**
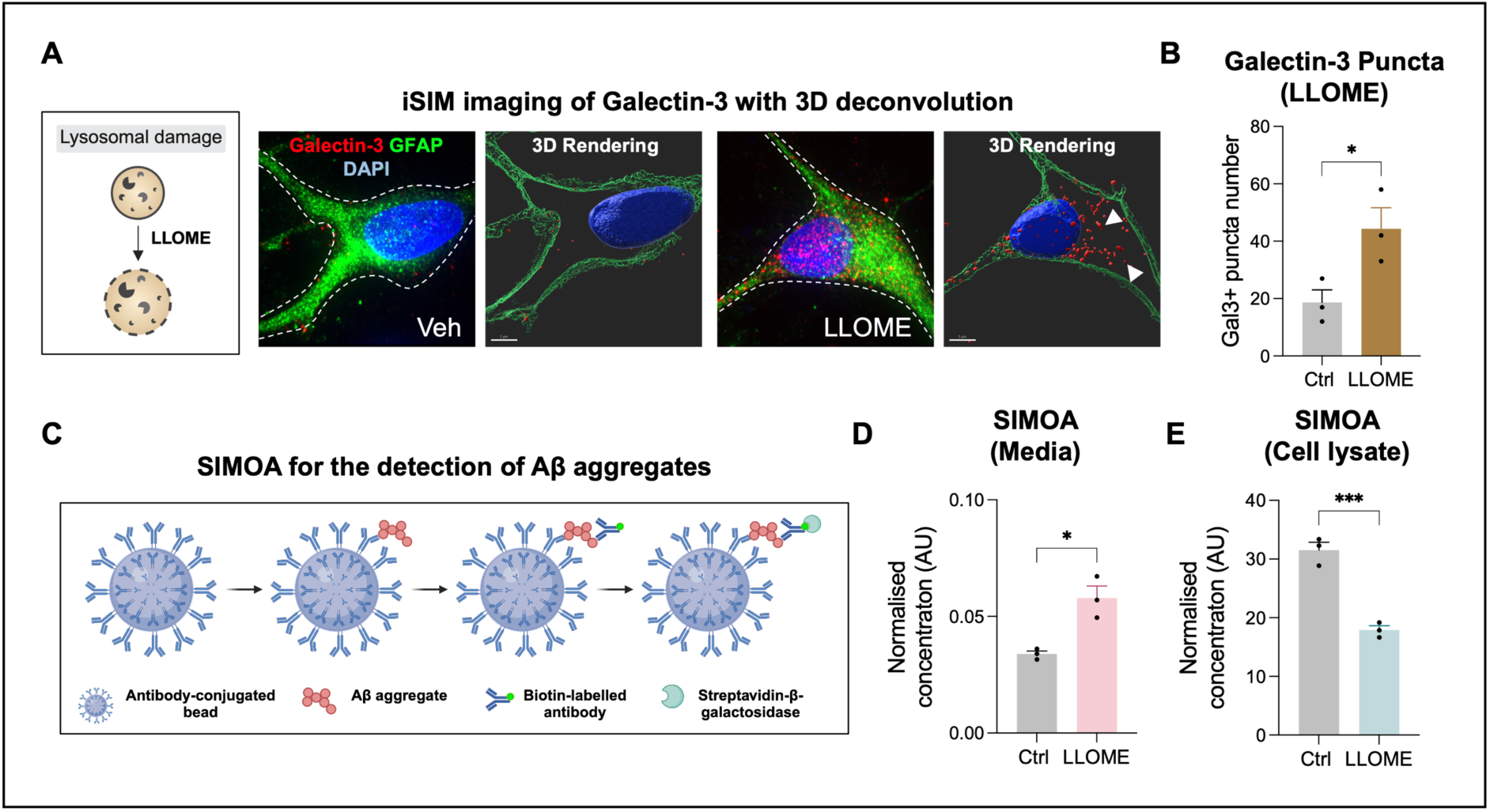
Lysosomal damage enhances extracellular Aβ aggregate release from human iPSC-derived astrocytes. (A) Schematic and iSIM images showing Galectin-3 puncta formation in astrocytes treated with the lysosomotropic agent L-leucyl-L-leucine methyl ester (LLOMe), indicative of lysosomal membrane damage. Scale bars = 5 µm. (B) Quantification of Galectin-3+ puncta confirm increased lysosomal damage in LLOMe-treated astrocytes. Data represent mean ± SEM from 3 biological replicates. (C) Schematic of the Simoa assay used for ultrasensitive detection of soluble Aβ aggregates. (D–E) Simoa reveals a significant increase in Aβ aggregates in conditioned media (D) and a reduction in the cell lysate (E) following LLOMe treatment. Data represent mean ± SEM from 3 biological replicates. *p < 0.05, **p < 0.01, ***p < 0.001 by unpaired t-test.

We then asked whether this lysosomal stress influences the release of Aβ aggregates using the highly sensitive single-molecule immunoassay, single molecule array (Simoa). The Simoa assay, a previously validated method^26^, employs antibody-coated beads to capture Aβ-containing aggregates, followed by incubation with a biotinylated detector antibody. This results in the formation of an immunocomplex with streptavidin β-galactosidase (SBG), which produces a fluorescent readout through resorufin β-d-galactopyranoside (RGP) (**Figure 2C**). The digital detection capability of Simoa allows precise quantification of low-abundance Aβ aggregates in the low picomolar range.

We observed a significant increase in secreted Aβ aggregates in the media of LLOMe-treated astrocytes (**Figure 2D**), coupled with a marked reduction of intracellular Aβ aggregates (**Figure 2E**), indicating that lysosomal membrane damage promotes the extracellular release of Aβ.

To confirm and expand on this finding, we performed DNA-PAINT super-resolution imaging of Aβ aggregates isolated from both media and lysates. LLOMe treatment led to a significant increase in the number of aggregates detected in the media (**Figure 3A**), while aggregates within the cell lysate were significantly reduced (**Figure 3B**). These data corroborate the Simoa results and indicate that lysosomal stress facilitates the release of astrocyte-derived Aβ aggregates.

**Figure 3.**
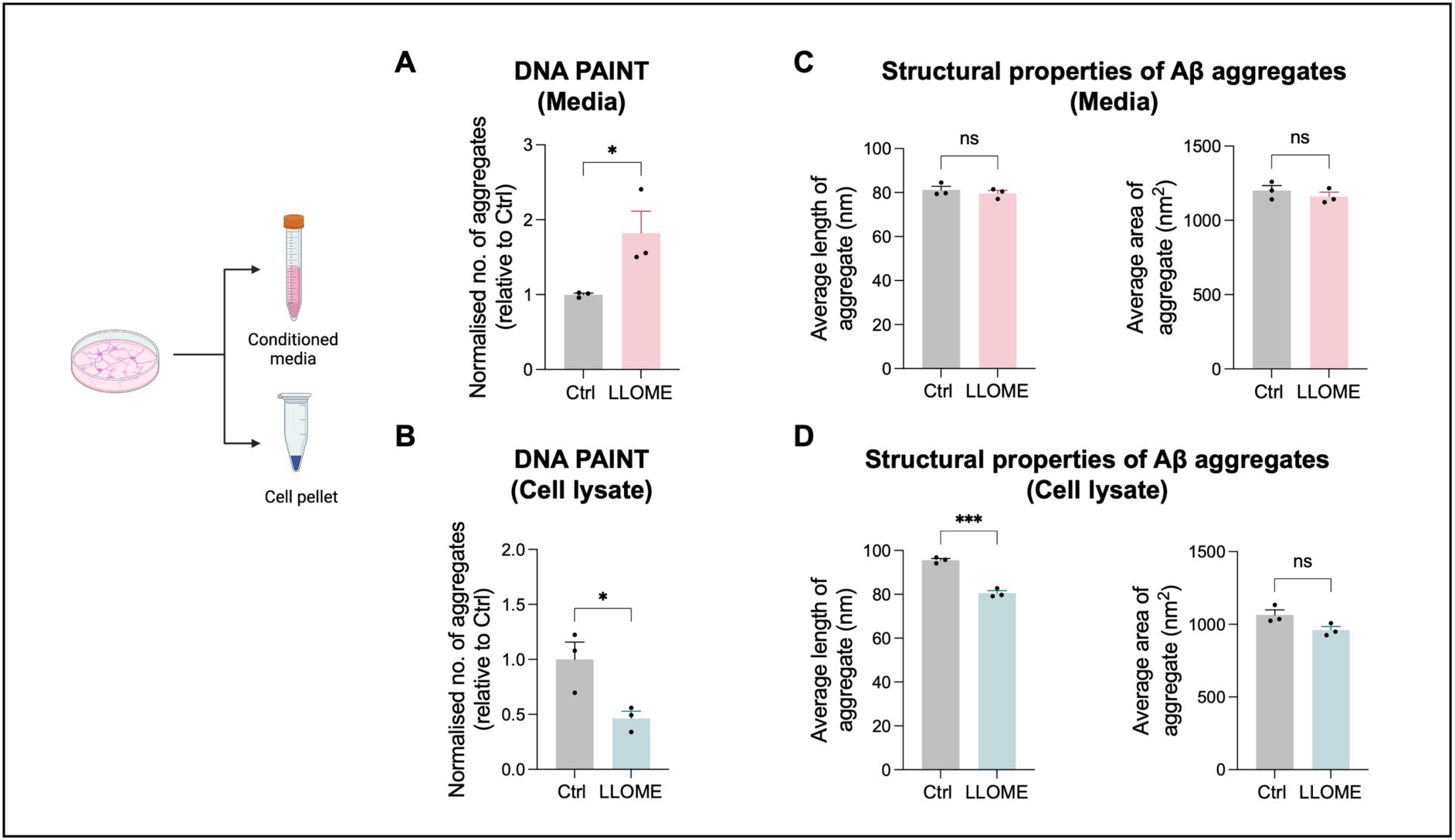
DNA-PAINT super-resolution imaging of Aβ aggregates isolated from both media and lysates. (A–B) DNA-PAINT confirms increased number of Aβ aggregates in the media (A) and a corresponding decrease in the lysate (B) after LLOMe. Data represent mean ± SEM from 3 biological replicates. (C–D) Structural analysis shows no change in aggregate length or area in the media (C), but a significant reduction in intracellular aggregate length in the lysate (D), with no difference in area Data represent mean ± SEM from 3 biological replicates. *p < 0.05, ***p < 0.001, ns = not significant by unpaired t-test.

We next examined whether lysosomal damage alters the structural characteristics of the released aggregates. In the media, the average length and area of Aβ aggregates remained unchanged following LLOMe treatment (**Figure 3C**). However, in the cell lysates, the average length of intracellular aggregates was significantly reduced, while their area remained unchanged (**Figure 3D**). These findings suggest that lysosomal damage promotes the release of aggregates without substantially altering their morphology in the extracellular environment.

Together, these results indicate that lysosomal membrane permeabilization triggers the release of Aβ aggregates from astrocytes into the extracellular space and may contribute to Aβ pathology under conditions of lysosomal stress.

### Lysosomal damage triggers changes in the transcriptome and secretome suggestive of reactive transformation in hiPSC-derived astrocytes

Having established that lysosomal stress promotes the extracellular release of Aβ aggregates from human astrocytes, we next sought to investigate whether this stress also triggers a reactive transformation. Reactive astrocytes are known to upregulate inflammatory genes and secrete cytokines and chemokines in response to cellular stress. To explore these molecular changes in detail, we performed transcriptomic and secretory profile analyses to define the astrocytic response to lysosomal membrane damage.

RNA-seq revealed significant upregulation of pro-inflammatory genes, suggesting a transition to a reactive state (**Figure 4A**). Key upregulated genes included *CCL2*, *CCL20*, *CCL5*, *ICAM1*, *SERPINE1*, *HBEGF*, *IL6*, and *IL8*, which are involved in immune response, cytokine signaling, and cellular stress responses. These findings indicate that LLOMe-treated astrocytes adopt a pro-inflammatory reactive state. Gene Set Enrichment Analysis (GSEA) of the RNA-seq data revealed significant enrichment of pathways associated with immune responses, chemokine signaling, and cellular stress, which are characteristic of reactive astrocyte transformation (**Figure 4B**). The most enriched pathway was “response to chemokine” (NES = 2.09, FDR < 0.001), highlighting the activation of genes involved in immune cell recruitment. Other enriched pathways included cytokine-mediated signaling (NES = 1.98, FDR < 0.001) and regulation of apoptotic process (NES = 1.92, FDR < 0.01), suggesting that LLOMe induces pro-inflammatory and stress-related responses in astrocytes.

**Figure 4.**
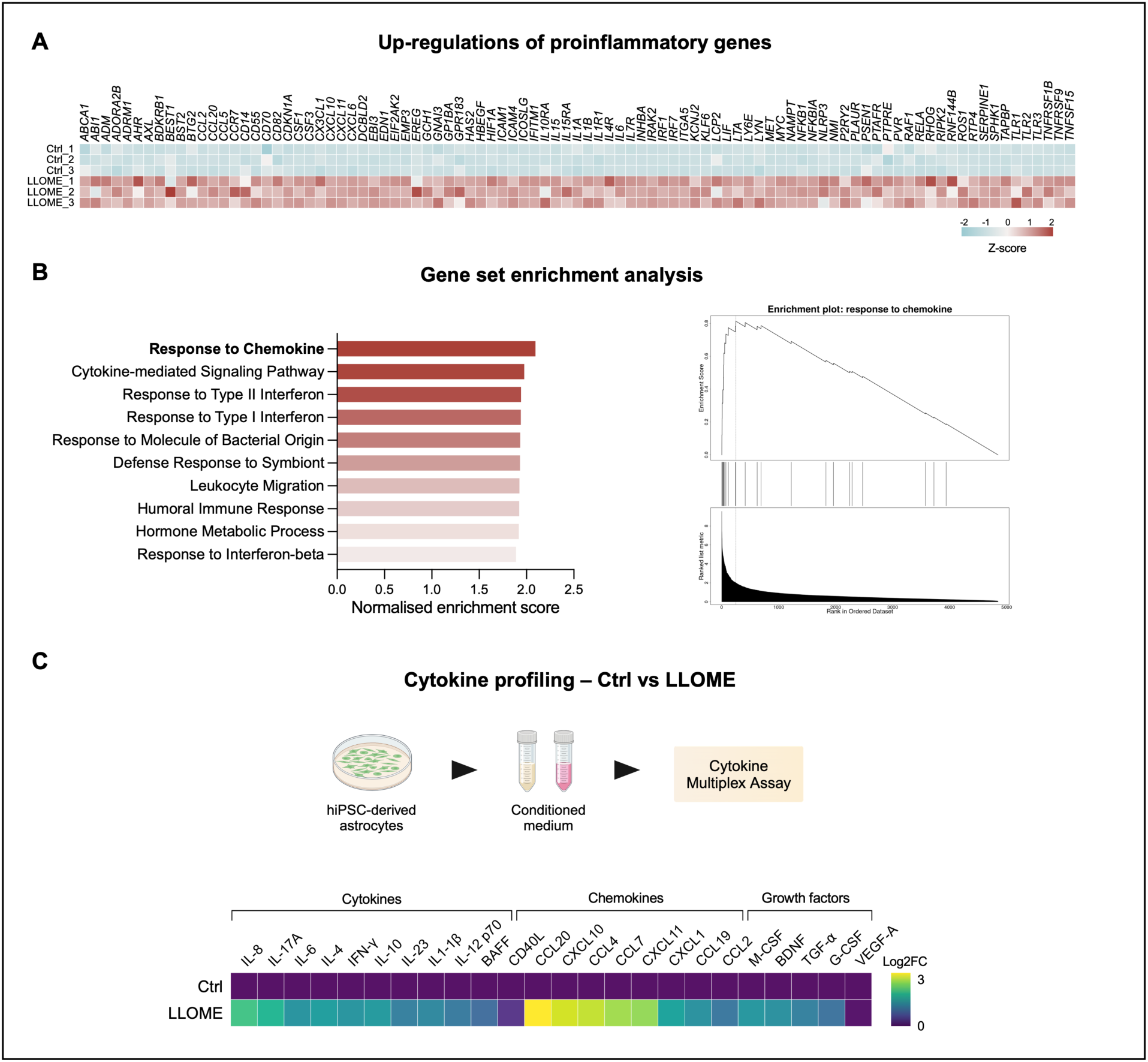
Functional characterization of LLOMe-induced cytokine and chemokine release. (A) RNA-seq analysis on the mRNA expression levels of proinflammatory genes. (B) Gene set enrichment analysis showing the fold enrichment based on the most up-regulated list of genes. (C) Schematic diagram of the experimental strategy (up). Multiplex immunoassay of cytokine and chemokine (bottom). Data represents 3 biological replicates.

To further characterize the functional impact of LLOMe treatment, we analyzed the release of cytokines, chemokines, metallopeptidases, and growth factors from the culture supernatant. This analysis revealed significant changes in the levels of key immune-related molecules, including IL-8, M-CSF, CXCL1, CCL2, IL-4, IL-17A, CXCL11, TGF-α, BDNF, CCL4, IL1-β, CCL7, IL-10, CCL20, VEGF-A, and CXCL10 (**Figure 4C**). These findings support the conclusion that lysosomal damage promotes a reactive transformation in human astrocytes characterized by a transcriptionally and functionally proinflammatory phenotype.

## Discussion

In this study, we demonstrate that human iPSC-derived astrocytes produce small, diffusible Aβ aggregates under basal conditions and that these aggregates are released into the extracellular space. Using DNA-PAINT super-resolution imaging, we characterized these structures in both cell lysates and conditioned media, revealing that the majority fall within the size range previously associated with synaptotoxicity in AD. Our findings challenge the prevailing view that Aβ aggregates originate primarily from neurons and support a growing body of evidence that glial cells, including astrocytes and oligodendrocytes, may directly contribute to Aβ pathology.

Importantly, we show that lysosomal membrane permeabilization via LLOMe markedly enhances the extracellular release of Aβ aggregates from astrocytes. This was accompanied by a decrease in intracellular aggregates, suggesting that lysosomal damage facilitates the redistribution of intracellular Aβ pools into the extracellular environment. Although the structural properties of the aggregates remained largely unchanged, their increased release may exacerbate extracellular Aβ burden and toxicity.

We further demonstrate that lysosomal stress drives a reactive transformation in astrocytes, characterized by transcriptional upregulation of pro-inflammatory genes and increased secretion of cytokines and chemokines. This dual response—release of potentially neurotoxic aggregates and induction of a reactive, inflammatory state— highlights lysosomal integrity as a central regulator of astrocyte behavior in AD-relevant contexts.

Our findings builds on recent studies implicating glial cells in Aβ pathology, including work showing that oligodendrocytes can generate Aβ and contribute to plaque formation in mouse models^2–4^. Together, these results broaden the understanding of cell-autonomous contributions to amyloid pathology and underscore the need to consider astrocytes not just as passive responders or clearance cells, but as active sources of Aβ under stress conditions.

Future studies are needed to explore the upstream triggers of astrocytic Aβ production, to validate these mechanisms *in vivo*, and to determine whether astrocyte-derived aggregates differ functionally or structurally from those of neuronal origin. Targeting lysosomal pathways in astrocytes may represent a promising therapeutic strategy to curb both amyloid accumulation and glial-mediated neuroinflammation in AD.

## Methods

### Reagents and Tools Table

| Reagent/Resource | Reference or Source | Identifier or Catalog Number |
| --- | --- | --- |
| **Experimental Models** |  |  |
| Human: CTRL1 iPSC line | This study | N/A |
| Human: CTRL2 iPSC line | This study | N/A |
| Human: CTRL3 iPSC line | This study | N/A |
| **Antibodies** |  |  |
| anti-mouse Galectin-3 | Biolegend | Cat#126702 |
| anti-chicken GFP-A | Abcam | Cat#Ab4674 |
| **Chemicals, Enzymes and other reagents** |  |  |
| Essential 8 Medium media | Life Technologies | Cat#A1517001 |
| Geltrex | Life Technologies | Cat#A1413302 |
| EDTA | Life Technologies | Cat#15575020 |
| Dorsomorphin | Tocris Bioscience | Cat#3093 |
| SB431542 | Tocris Bioscience | Cat#1614 |
| RiboLock RNase Inhibitor | Thermo Fisher Scientific | Cat#EO0381 |
| Formamide | Thermo Fisher Scientific | Cat#17899 |
| UltraPure BSA | Thermo Fisher Scientific | Cat#AM2616 |
| **Software** |  |  |
| R project for Statistical Computing v4.2.0 | CRAN | RRID: SCR_001905 |
| DESeq2 | Bioconductor | RRID: SCR_015687 |
| Salmon | https://combine-lab.github.io/salmon/ | RRID: SCR_017036 |

### Methods and Protocols

#### Differentiation of hiPSC-derived astrocytes

Human iPSCs were maintained on Geltrex (Life Technologies) with Essential 8 Medium media (Life Technologies) and passaged using EDTA. Generation of hiPSC-derived astrocytes was carried out based on a previously described protocol^20^. Briefly, after neural conversion (12 days in a 1:1 mixture of N2- and B27-containing media containing 1 μM Dorsomorphin (Millipore) and 10 μM SB431542 (Tocris Bioscience)). After a propagation phase (>60 days) with 10 ng/ml FGF-2 (Peprotech) and were terminally differentiated to astrocytes using BMP4 (10 ng/ml, R&D) and LIF (10 ng/ml, Sigma-Aldrich).

#### Image acquisition and analysis

For immunostaining, cells were fixed in 4% paraformaldehyde (PFA) in PBS followed by permeabilization with Triton X-100 (Sigma). Fixed cells were blocked with 10% normal goat serum (Sigma) in PBS, probed with primary antibodies (anti-mouse Galectin-3 (BioLegend) and anti-chicken GFPA (Abcam)) diluted in blocking solution and detected with goat anti-mouse or anti-chicken secondary antibodies coupled to Alexa Fluor 488 or 594.

For the analysis of vesicle size, fixed cells were imaged on an iSIM microscope using a 150× oil objective with deconvolution. Galectin 3+ puncta quantifications were done using the spot detection function in the Imaris software. Surface masks were created using the GFAP staining to indicate astrocytes.

#### Protein analysis

Extracellular cytokine, chemokine, metallopeptidase, and growth factor were measured in conditioned media using Proteome Profiler Human XL Cytokine Array Kit (Bio-Techne Ltd.) according to the manufacturer’s instructions. Conditioned media samples from experiments collected at different time points were frozen at −80°C.

#### RNA-seq analysis

Sequencing data was processed using nf-core/rnaseq ^27^, which is part of the nf-core collection of workflows. Reads underwent pre-processing, including quality filtering and trimming, before being mapped to the GRCh37 genome. The tools and software used during this process included FastQC (version 0.11.9) for quality control, Cutadapt (version 0.6.7) and Trim Galore (version 3.4) for trimming, and Salmon (version 1.10.1) for quantification. Custom scripts were used for various steps, written in Python (version 3.9.5) and YAML (version 6.0).

The mapping and alignment of reads were performed using STAR (version 2.7.10a), and Samtools (version 1.10) was used for post-alignment processing. Additionally, UMI-tools (version 1.1.4) was used for deduplication, and Bioconductor’s tximeta (version 1.12.0) and Summarized Experiment (version 1.24.0) packages were employed for transcript quantification and gene expression analysis. All workflows were executed using Nextflow (version 22.10.7) with nf-core/rnaseq (version 3.12.0).

Mapped reads were de-duplicated and quantified to generate gene counts for differential expression analysis. Differential gene expression was analyzed using the SARTools pipeline ^28^, and statistical analysis was performed with DESeq2 ^29^, version 1.40.2.

#### Quantification of soluble Aβ aggregates

Single-molecule pull-down assays were conducted following previously established protocols^30^. In brief, glass coverslips coated with polyethylene glycol (PEG) and biotin were treated with neutravidin (0.2 mg/ml) in TBS containing 0.05% Tween 20 (TBST) for 10 minutes. The coverslips were then washed twice with TBST and once with TBS containing 1% Tween 20 (1%T).

Next, biotinylated 6E10 antibody (10 nM; BioLegend; 803007) was applied for 15 minutes, followed by two TBST washes and one with 1%T. Media samples were then added and incubated overnight at 4°C, followed by two additional TBST washes and one with 1%T. The coverslips were then incubated with a secondary 6E10 antibody (500 pM; BioLegend; 803020) conjugated to single-strand DNA (ACCACCA) for 45 minutes, followed by two washes with TBST and one with 1%T.

After washing, TetraSpeck microspheres (1:7,000 dilution in TBS, 10 μl; Thermo Scientific, Cat. T7279) were introduced to each well for 10 minutes. The TetraSpeck solution was subsequently removed, followed by two additional TBST washes. A second PDMS gasket (Merck, GBL-103250-10EA) was stacked onto the coverslip before adding 3 μl of a complementary imaging strand (TGGTGGT-Cy3B; atdbio) in TBS. Finally, the coverslip was sealed by placing another coverslip over the second PDMS gasket.

Images were captured using a custom-built TIRF microscope^30^ equipped with a 520 nm laser, with an exposure time of 100 milliseconds for 4,000 frames per field of view. Three fields of view were acquired per well. The acquired frames were stacked, reconstructed, drift-corrected, and analyzed to quantify aggregate number and length using ACT software^31^. Images of individual aggregates were further processed using the ASAP software^32^.

#### Simoa Plate Preparation

Antibody-bead conjugation was carried out according to the Quanterix Homebrew instruction manual. Simoa plates were prepared using a “3-step assay” format, following the Quanterix protocol. The final plate was processed using the Quanterix SR-X Instrument. To maintain a digital readout, we ensured that no sample or calibrant concentrations exceeded an fON value of 0.7, preventing the transition into an analog detection mode where signal intensity is influenced by bead brightness rather than bead count.

#### Quantification and statistical analysis

Unless otherwise specified, statistics analysis was performed using GraphPad Prism (Version 10). Student’s t test was used to compare differences between two groups. One-way ANOVA followed by post testing with Dunnett’s method was used to analysis differences between more than two groups. For precise p value calculation, a multiple t test was performed after ANOVA calculations. Significance threshold was defined as adjusted p value < 0.05. Error bars in all figures represent SEM. The number of replicates (n) is listed in the legend of each figure.

## Declarations

### Ethics approval and consent to participate

Not applicable.

### Consent for publication

Not applicable.

### Availability of data and materials

Materials generated in this study may be available from the Lead Contact with a completed Materials Transfer Agreement. Restrictions may apply to the availability of the cell lines due to our need to maintain the stock. The raw FASTQ files from the RNA-seq analysis have been deposited in the GEO (Gene Expression Omnibus) database: GSE279326.

### Authors’ contributions

C.H. and R.P. conceived and designed the study. E.F. and D.K. conceived and designed the experiments on single-molecule immunoassays. F.J.L. contributed cell lines. S.G. performed the data analysis on the RNA sequencing datasets. P.D.B. prepared the RNA library, and S.D. performed library QC and sequencing. All authors read and approved the manuscript.

## Acknowledgements

The authors wish to thank the patients for fibroblast donation and UCL Genomics facility for performing RNA-seq analysis.

## References

1. Selkoe, D. J. & Hardy, J. The amyloid hypothesis of Alzheimer’s disease at 25 years. EMBO Mol. Med. 8, 595–608 (2016).

2. Sasmita, A. O. et al. Oligodendrocytes produce amyloid-β and contribute to plaque formation alongside neurons in Alzheimer’s disease model mice. Nat. Neurosci. 27, 1668–1674 (2024).

3. Ishii, A. et al. Contribution of amyloid deposition from oligodendrocytes in a mouse model of Alzheimer’s disease. Mol. Neurodegener. 19, 83 (2024).

4. Rajani, R. M. et al. Selective suppression of oligodendrocyte-derived amyloid beta rescues neuronal dysfunction in Alzheimer’s disease. PLoS Biol. 22, e3002727 (2024).

5. Thal, D. R. The role of astrocytes in amyloid β-protein toxicity and clearance. Exp. Neurol. 236, 1–5 (2012).

6. Kim, J., Yoo, I. D., Lim, J. & Moon, J.-S. Pathological phenotypes of astrocytes in Alzheimer’s disease. Exp. Mol. Med. 1–5 (2024).

7. Patani, R., Hardingham, G. E. & Liddelow, S. A. Functional roles of reactive astrocytes in neuroinflammation and neurodegeneration. Nat. Rev. Neurol. 1–15 (2023).

8. Brandebura, A. N., Paumier, A., Onur, T. S. & Allen, N. J. Astrocyte contribution to dysfunction, risk and progression in neurodegenerative disorders. Nat. Rev. Neurosci. 24, 23–39 (2023).

9. Liddelow, S. A. et al. Neurotoxic reactive astrocytes are induced by activated microglia. Nature 541, 481–487 (2017).

10. Di Malta, C., Fryer, J. D., Settembre, C. & Ballabio, A. Astrocyte dysfunction triggers neurodegeneration in a lysosomal storage disorder. Proc. Natl. Acad. Sci. 109, E2334–E2342 (2012).

11. Hung, C. & Livesey, F. J. Endolysosome and autophagy dysfunction in Alzheimer disease. Autophagy 17, 3882–3883 (2021).

12. Hung, C. O. Y. & Livesey, F. J. Altered γ-Secretase Processing of APP Disrupts Lysosome and Autophagosome Function in Monogenic Alzheimer’s Disease. Cell Rep. 25, 3647–3660 (2018).

13. Hung, C. et al. SORL1 deficiency in human excitatory neurons causes APP-dependent defects in the endolysosome-autophagy network. Cell Rep. 35, 109259 (2021).

14. Nixon, R. A. The role of autophagy in neurodegenerative disease. Nat. Med. 19, 983–997 (2013).

15. Colacurcio, D. J., Pensalfini, A., Jiang, Y. & Nixon, R. A. Dysfunction of autophagy and endosomal-lysosomal pathways: Roles in pathogenesis of Down syndrome and Alzheimer’s Disease. Free Radic. Biol. Med. 114, 40–51 (2018).

16. Nixon, R. A. Amyloid precursor protein and endosomal–lysosomal dysfunction in Alzheimer’s disease: inseparable partners in a multifactorial disease. FASEB J. 31, 2729–2743 (2017).

17. Jiang, Y. et al. Lysosomal dysfunction in Down syndrome is APP-dependent and mediated by APP-βCTF (C99). J. Neurosci. 39, 5255–5268 (2019).

18. Papadopoulos, C., Kravic, B. & Meyer, H. Repair or lysophagy: dealing with damaged lysosomes. J. Mol. Biol. 432, 231–239 (2020).

19. Wang, F., Gómez-Sintes, R. & Boya, P. Lysosomal membrane permeabilization and cell death. Traffic 19, 918–931 (2018).

20. Taha, D. M. et al. Astrocytes display cell autonomous and diverse early reactive states in familial amyotrophic lateral sclerosis. Brain 145, 481–489 (2022).

21. Whiten, D. R. et al. Nanoscopic characterisation of individual endogenous protein aggregates in human neuronal cells. ChemBioChem 19, 2033–2038 (2018).

22. Fertan, E. et al. Cerebral organoids with chromosome 21 trisomy secrete Alzheimer’s disease-related soluble aggregates detectable by single-molecule-fluorescence and super-resolution microscopy. Mol. Psychiatry 29, 369–386 (2024).

23. Fertan, E. et al. Clearance of beta-amyloid and tau aggregates is size dependent and altered by an inflammatory challenge. Brain Commun. 7, fcae454 (2025).

24. Hung, C., Fertan, E., Livesey, F. J., Klenerman, D. & Patani, R. APP antisense oligonucleotides reduce amyloid-β aggregation and rescue endolysosomal dysfunction in Alzheimer’s disease. Brain 147, 2325–2333 (2024).

25. Stern, A. M. et al. Abundant Aβ fibrils in ultracentrifugal supernatants of aqueous extracts from Alzheimer’s disease brains. Neuron 111, 2012–2020 (2023).

26. Böken, D. et al. Ultrasensitive Protein Aggregate Quantification Assays for Neurodegenerative Diseases on the Simoa Platform. Anal. Chem. 97, 290–299 (2024).

27. Patel, H. et al. nf-core/rnaseq: nf-core/rnaseq v3. 12.0-Osmium Octopus. (2023).

28. Varet, H., Brillet-Guéguen, L., Coppée, J.-Y. & Dillies, M.-A. SARTools: a DESeq2-and EdgeR-based R pipeline for comprehensive differential analysis of RNA-Seq data. PLoS One 11, e0157022 (2016).

29. Love, M. I., Huber, W. & Anders, S. Moderated estimation of fold change and dispersion for RNA-seq data with DESeq2. Genome Biol. 15, 1–21 (2014).

30. Emin, D. et al. Small soluble α-synuclein aggregates are the toxic species in Parkinson’s disease. Nat. Commun. 13, 5512 (2022).

31. Xia, Z., et al. A computational suite for the structural and functional characterization of amyloid aggregates. Cell Reports Methods 3, (2023).

32. Danial, J. S. H. & Garcia-Saez, A. J. Quantitative analysis of super-resolved structures using ASAP. Nat. Methods 16, 711–714 (2019).

